# Long-Term Imaging of Living Adult Zebrafish

**DOI:** 10.1101/2021.03.19.436170

**Authors:** Daniel Castranova, Bakary Samasa, Marina Venero Galanternik, Aniket V. Gore, Brant M. Weinstein

## Abstract

The zebrafish has become a widely used animal model due in large part to its accessibility to and usefulness for high-resolution optical imaging. Although zebrafish research has historically focused mostly on early development, in recent years the fish has increasingly been used to study regeneration, cancer metastasis, behavior, and other processes taking place in juvenile and adult animals. However, imaging of live adult zebrafish is extremely challenging, with survival of adult fish limited to a few tens of minutes using standard imaging methods developed for zebrafish embryos and larvae. Here, we describe a new method for imaging intubated adult zebrafish using a specially designed 3D printed chamber for long-term imaging of adult zebrafish on inverted microscope systems. We demonstrate the utility of this new system by nearly day-long observation of neutrophil recruitment to a wound area in living double-transgenic adult casper zebrafish with fluorescently labeled neutrophils and lymphatic vessels.

## INTRODUCTION

The zebrafish is an excellent animal model to study a variety of biological processes. Like humans, zebrafish are vertebrates, and fish share many of our genes and develop many of the same sorts of diseases and pathologies that affect us. However, they have a variety of useful practical advantages for genetic and experimental studies. A two day post fertilization embryo is just 3 mm long and an adult is only about 2-4 cm in length, which allows for many fish to be housed in a relatively small space. Zebrafish are robust, developing rapidly from embryo to adult in 2 to 3 months. They can breed every week or two and produce around 200 embryos per spawning event, providing many animals for experimental analysis. Zebrafish embryos are externally fertilized and all stages of their development are readily available for observation and experimental manipulation. Finally, and perhaps most importantly, the optical clarity of zebrafish embryos and early larvae makes it possible to carry out very high-resolution optical imaging of all developing organs and tissues, including those deep within the animal.

These practical advantages have been amplified and extended upon by a variety of tools and methods developed for the fish. These include ENU mutagenesis screens that have identified thousands of mutations involved in embryonic development (Driever et al., 1996; Mullins et al., 1994), the subsequent characterization of which has led to innumerable important new insights (Patton and Zon, 2001). Reverse genetic tools such as morpholinos (Nasevicius and Ekker, 2000) and CRISPR (Hwang et al., 2013) can also be used to target specific genes of interest in fish. A wide variety of different transgenic lines have also been developed for spatial and temporal control of gene expression in fish (Scheer and Campos-Ortega, 1999; Wyart et al., 2009) permitting functional manipulation of the entire organism, specific tissues, or even single cells. Tens of thousands of zebrafish lines have been created (zfin.org), expressing fluorescent proteins in different organs and tissues, facilitating detailed observation and study of tissue and organ development using fluorescence microscopy. Use of these transgenic reporter lines has led to advances in understanding the function of many organs and tissues, including blood and lymphatic vessels (Jung et al., 2017; Lawson and Weinstein, 2002; Yaniv et al., 2006), the nervous system (Wyart and Del Bene, 2011), the liver and pancreas (Ober et al., 2003), and many others.

Although the historical focus of zebrafish research has been on embryonic and larval development, there has recently been increasing interest in using zebrafish to study processes such as disease and tissue regeneration in adults. Research using adult zebrafish has already led to novel insights into cancer metastasis (Kaufman et al., 2016) with great potential for developing new treatments (Frantz and Ceol, 2020). Human tumors transplanted into zebrafish are being used as a platform for developing precision cancer therapies (Fazio et al., 2020; Yan et al., 2019). Zebrafish have also been used to model obesity and diabetes, with recent work showing that the pathophysiological pathways are conserved between mammals and zebrafish (Oka et al., 2010). The ability of zebrafish to regenerate complex tissues including the fins (Johnson and Weston, 1995) and heart (Poss et al., 2002), makes them a superb model for understanding mechanisms controlling tissue regeneration. Furthermore, there are tissues that do not exist during early embryonic and larval development such as the newly described intracranial lymphatic vascular network (Castranova et al., 2021) that can only be imaged and studied in juvenile and adult stage fish.

While there has been significant progress in the use of adult zebrafish as a research model, microscopic imaging of adults remains extremely challenging. Even if they are kept submerged, anesthetized adult zebrafish can only be imaged for a few tens of minutes before the animal dies, generally from lack of adequate oxygenation. In terrestrial models, longer-term live imaging generally involves anesthesia with intubation to ensure that the animal remains oxygenated and viable during imaging. A previous report from Xu et al. described a method for intubating and holding adult zebrafish for live imaging on an upright microscope (Xu et al., 2015). Here, we describe a new and further refined method designed for inverted microscopes, providing a detailed protocol and design plans that incorporate a variety of useful and important new additions, such as a multiple safety features to prevent water overflow into objective lenses and microscope hardware. We provide easy-to-use instructions for replicating this method including downloadable CAD designs for imaging chambers that can be easily adapted to any microscope system and used to create chambers with widely available 3-D thermoplastic printers. We demonstrate the power and utility of our system by carrying out long-term imaging of neutrophil recruitment to a wound in an adult zebrafish with transgenically labeled neutrophils and lymphatic vessels.

## MATERIALS AND METHODS

### Construction of the Intubation and Imaging Chamber

A 3D printable plastic chamber was designed that would hold a glass-bottomed chambered coverglass, allow inflow and outflow tubing to enter and exit, hold a water sensor and an overflow tube, and fit on our existing Tokei Hit heated microscope stage (Model: INUB-TIZB; **Fig 1E**). A second version of the chamber was also designed to fit on any stage designed to accept a 96 well plate (example: Tokei Hit Model: STZF-TIZWX-SET), with four supporting arms which can be quickly adjusted in CAD software to fit any other microscope stage (**Fig 1F**). 3D models for the printable plastic chambers were designed using Google SketchUp Pro and these CAD designed are included as supplements (**intubation_chamber**.**stl, 96W_Intubation_chamber**.**stl**). They can be downloaded and used to print the imaging chambers described above. These CAD designs are easily be modified to create chambers to fit other microscope stages. Chambers can be printed in-house if 3D printing is available, or there are many “print and ship” options available. The material used for printing must be watertight and inert. Our chambers were printed out of Nylon 12 by XometryTM (xometry.com) using Selective Laser Sintelating (SLS).

**Figure 1.**
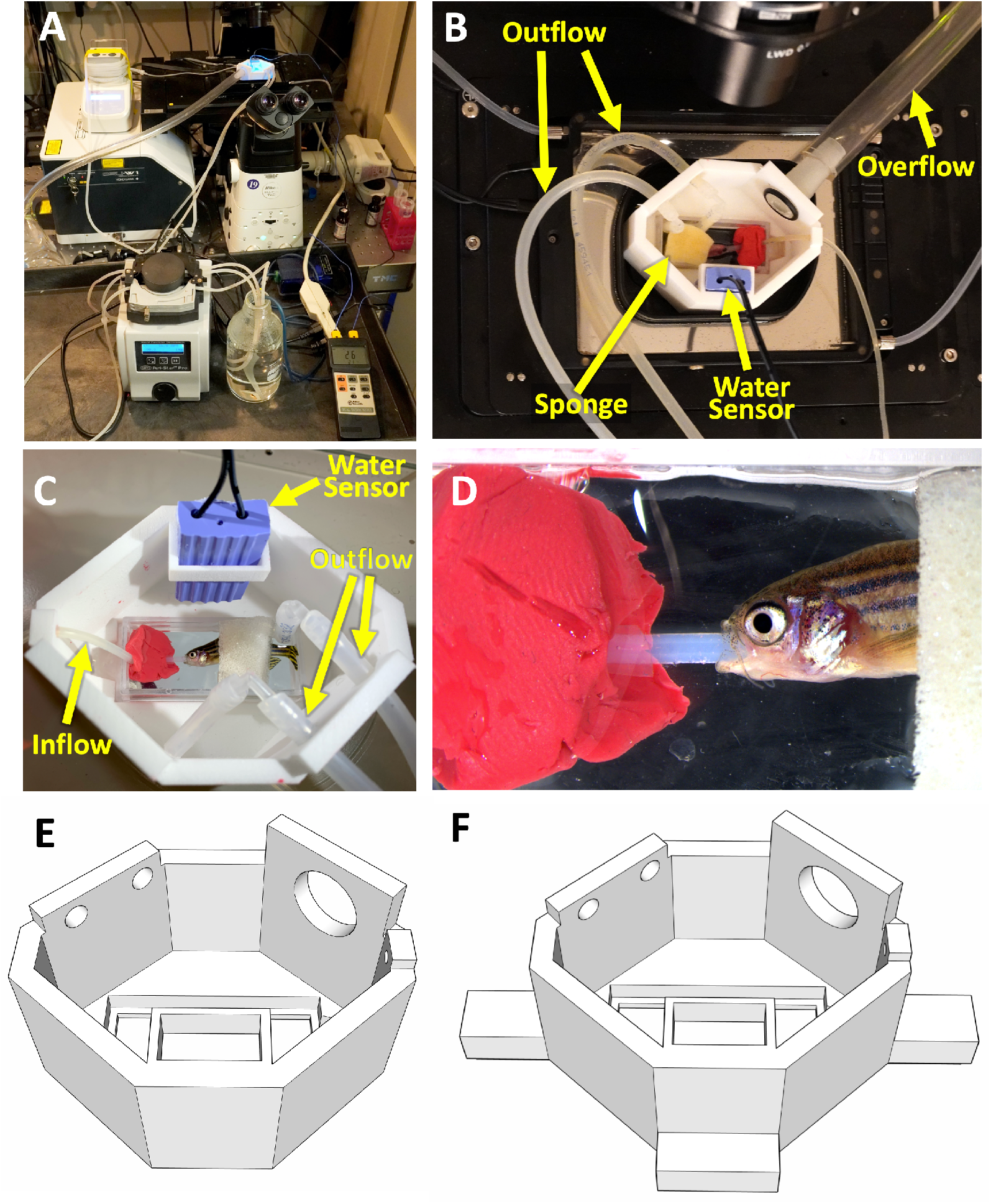
Intubation of an adult fish on an inverted microscope. **A-D**. Photographs of long-term imaging of an Intubated adult zebrafish inside custom 3D printed chamber. **A**. Overview photograph of an intubation rig set up on an inverted microscope. **B**. Photograph of the intubation/imaging chamber set up on a microscope stage, showing inflow tube, dual outflow tubes, overflow tube, water sensor, and sponge holding fish in place. **C**. Close-up photograph of the chamber with an intubated fish, with inflow tube, dual outflow tubes, and water sensor noted. **D**. Higher magnification image of the intubated fish. **E**,**F**. 3D renderings of intubation chambers designed to fit on (**E**) a Tokai Hit heated stage (Model: INUB-TIZB) or (**F**) in a 96 well plate holder on a Tokei Hit heated stage (Model: STZF-TIZWX-SET) or any other stage designed to hold 96 well plates.

A single-well chambered coverglass (Lab-TekII #155360) is placed into the opening in the bottom of the 3D printed chamber by placing a small bead of silicone grease (SG-ONETM 24708) around the outer edge of the chambered coverglass using a 12cc syringe and pressing it firmly in place (**Fig 2D**). The 3D printed chamber should be filled with water to test for leaks in the seal before use. Care should be taken to use only enough grease to create a water-tight seal. Excess grease can get on the bottom of the coverglass or worse, on the objective lens of the microscope.

**Figure 2.**
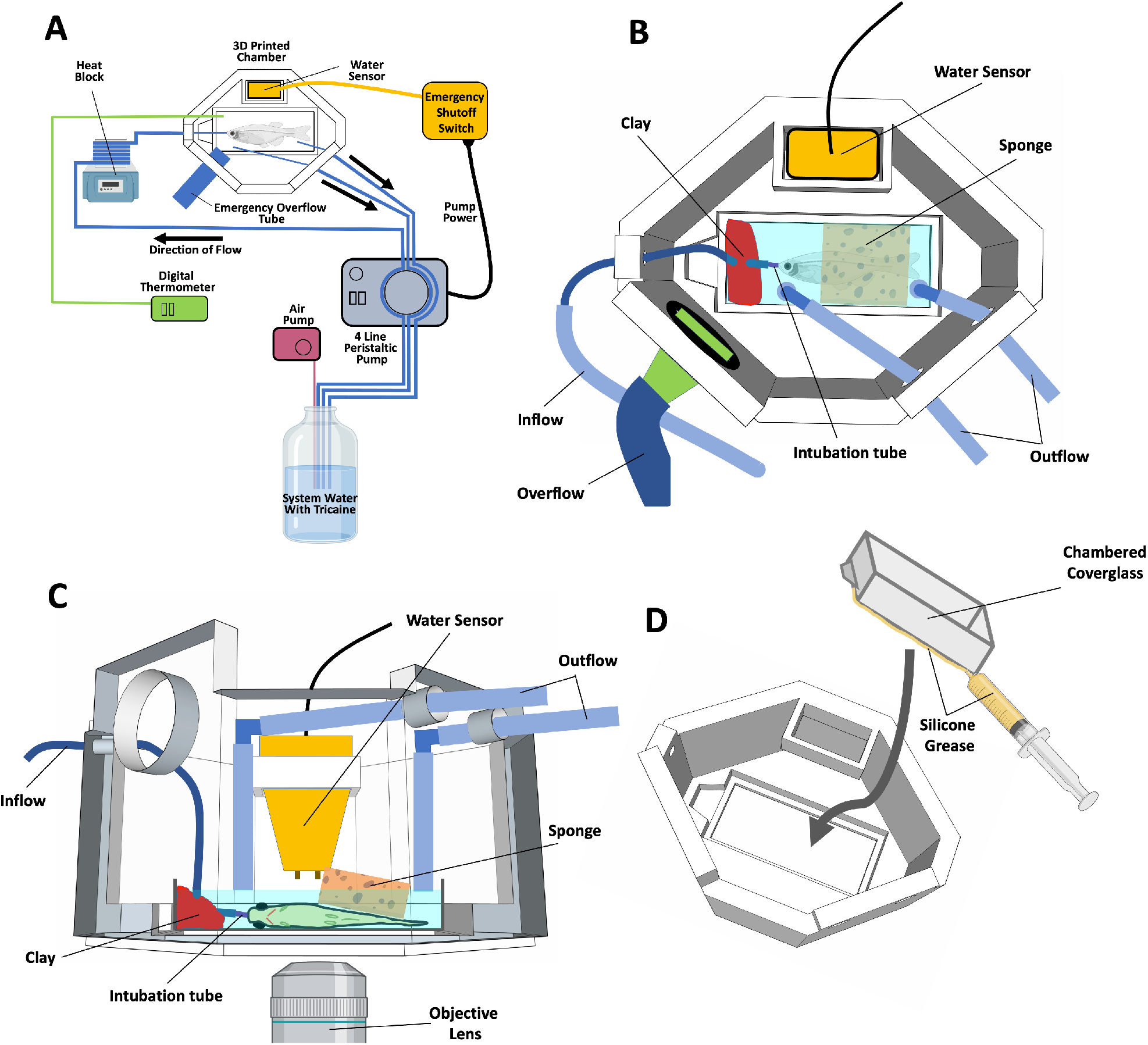
Schematic diagrams of zebrafish intubation. **A**. Schematic diagram providing an overview of the arrangement and interconnection of components for fish intubation. **B**. Schematic diagram showing a top view of the intubation chamber. **C**. Schematic diagram showing a side view of the intubation chamber. **D**. Schematic diagram showing attachment of a chambered coverglass into the 3D printed chamber, using silicone grease to provide a watertight seal.

### Water Flow

One liter of aquarium system water with 126 mg/L buffered tricaine in is placed in a bottle with an air stone inside to aerate the system water. A four-channel peristaltic pump (World Precision Instruments Peri-StarTM Pro Cat# PERIPRO −4L) is used to pump the aquarium system water through silicone tubing (Tygon) which is wrapped around a heat block (SH100 Mini Dry Bath Hot Block) to warm the water before it goes through the inflow hole in the 3D printed chamber (**Fig 2A**). Wrapping the tygon tubing around the heat block six times and setting the heat block to 57°C produced a water temperature between 26 and 28°C inside the chambered coverglass. The tube is held to the bottom of the chambered coverglass using modeling clay (**Fig 2B,C**). A small piece of tubing is placed in the end of the outflow tube, small enough to fit in the fish’s mouth (**Fig 2C**). Setting the peristaltic pump at 15 rotations per minute provides a flow rate of 6 ml per minute through the chambered coverglass. Two outflow tubes are connected to the same peristaltic pump and draw water out of the chambered coverglass and back into the bottle (**Fig 2A**). These tubes go through the two specially designed holes in the 3D printed chamber and attach to elbow fittings and another small length of tube (**Fig 2B,C**). The height that these outflow tubes are placed will determine the water level inside the chambered coverglass and should be set so that the fish is completely submerged but the chambered coverglass does not overflow (**Fig 2C**). The temperature of the water flowing through the imaging chamber was measured using a digital thermometer (Fisher Scientific Dual Thermometer 15-077-26) and maintained between 26-28° C by adjusting the temperature of the heat block.

### Overflow Prevention Safety Features

Overflow of water onto the microscope would be catastrophic, so we developed several features to prevent it. First, there is one small diameter tube (in the fish’s mouth) bringing water into the chambered coverglass but two redundant larger diameter tubes removing water so even if one tube becomes clogged the inflow rate will still not exceed the ouflow rate. Second, we incorporated an automatic electronic water overflow cut-off switch. We placed a WasherWatcher Laundry Tub Overflow Protector (Overflow Protector) (HydroCheck, STAK Enterprises Inc.) into a sensor holder designed into the 3D printed chamber (cut to fit) (**Fig. 1B,C**). The Overflow Protector is plugged into an outlet, and the peristaltic pump is plugged into the Overflow Protector. In the event water overflows the chambered coverglass and enters the 3D printed chamber, the water sensor will detect it and cut the power to the peristaltic pump, preventing additional water entering the chamber. Third, we designed a last-resort emergency overflow outlet from the chamber to prevent water overflowing the chamber if all else fails. A large hole in the upper part of the 3D printed chamber (**Fig. 1E,F**) accepts a fitting held in place by a greased (silicone grease) “O” ring attached to a large-diameter tube leading to an empty 1L bottle (**Fig. 1B** and **Fig. 2B**). If the other leak prevention measures fail and water reaches the large overflow hole, water will be drained safely away from the microscope and into the empty bottle.

### Fish Preparation

Adult *casper* double transgenic zebrafish, *Tg(lyz:DsRed2)NZ50;Tg(mrc1a:eGFP)y251*, not fed on the day of the intubation, were anesthetized in 168 mg/L buffered tricaine (note 25% higher than what circulates in the chamber) in system water. A small wound was made around the trunk mid-line above the anal fin using a 4 mm dissecting knife and a few scales were removed (Fine Science Tools # 10055-12). The fish was then placed into the chambered coverglass inside the 3D printed chamber with the wound side facing down. The chamber was placed on the stage of a nearby stereo microscope, then using two pairs of blunt forceps, one to manipulate the intubation tube and one to manipulate the fish, the intubation tube was carefully placed into the fish’s mouth. Turning off the peristaltic pump is helpful while placing the tube inside the fish’s mouth. To prevent movement during image acquisition a sponge cut to fit inside the chambered coverglass was gently placed on top of the fish (**Fig 2B,C**). If additional stabilization for higher magnification imaging is needed, stabilizing weights were used instead of a sponge. Stabilizing weights were made by cutting a finger off of a nitrile glove, filling it with glass beads (425-600 um Sigma G8772) and tying it off so the beads do not escape. If the fish being intubated needs to be revivied after intubation, we found that replacing the tricaine water with system water and letting it run through the rig for 10 to 15 minutes until the fish begins to move was the best way to recover fish.

### Fish Husbandry and Fish Strains

Fish were housed in a large zebrafish dedicated recirculating aquaculture facility (4 separate 22,000L systems) in 6L and 1.8L tanks. Fry were fed rotifers and adults were fed Gemma Micro 300 (Skretting) once per day. Water quality parameters were routinely measured and appropriate measures were taken to maintain water quality stability (water quality data available upon request). The following transgenic fish lines were used for this study: *Tg(mrc1a:eGFP*)*y251* (Jung et al., 2017), and *Tg(lyz:DsRed2)nz50* (Hall et al., 2007). Fish were maintained and imaged in a *casper* (*roy, nacre* double mutant (White et al., 2008)) genetic background in order to increase clarity for visualization by eliminating melanocyte and iridophore cell populations from distorting images. This study was performed in an AAALAC accredited facility under an active research project overseen by the NICHD ACUC, Animal Study Proposal # 18-015.

### Image Acquisition

Confocal images of lymphatics and neutrophils were acquired using a Nikon Ti2 inverted microscope with Yokogawa CSU-W1 spinning disk confocal, Hamamatsu Orca Flash 4 v3 camera with the following Nikon objectives: 4X Air 0.2 N.A., 10X Air 0.45 N.A., 20X Air 0.7 N.A. Stereo microscope pictures were taken using a Leica M165 microscope with Leica DFC 7000T camera. In addition to acquiring eGFP expressing lymphatics with a 488 nm excitation laser and DSred2 with a 561 nm laser, we also acquired autofluorescence in the scales using 405 nm excitation laser and used these images to delineate the wound area. Video of the chamber on the inverted microscope, and fish swimming (**Supp Movie 1**, 5-10”, and Supp Movie 2) were taken with an Iphone XR.

### Image Processing

Images were processed using Nikon Elements. Maximum intensity projections of confocal stacks are shown. Time-lapse movies were made using Nikon Elements and exported to Adobe Premiere Pro CC 2019. Adobe Premiere Pro CC 2019 and Adobe Photoshop CC 2019 were used to add labels and arrows to movies. Schematics were made using Adobe Photoshop CC 2019, Microsoft PowerPoint, and Bio Render software.

### Intubation system maintenance

After each use the intubation system should be cleaned by running a 1:10 bleach solution through the system for at least 10 minutes followed by at least two flushes of tap water to remove any residual bleach. The tube sections that are used in the peristaltic pump become worn out quickly and should be replaced between uses. All of the tubing needs to be replaced occasionally, especially if any mold, mildew, or bacterial growth is seen. The chambered coverglass can be cleaned with 70% ethanol between uses but also needs to be replaced when it can no longer be effectively cleaned.

## RESULTS AND DISCUSSION

Previously, we reported methods for short-term and long-term imaging of developing zebrafish embryos and larvae (Kamei and Weinstein, 2005), but these methods are not suitable for imaging adult zebrafish. We have now designed a modified, improved chamber and associated “rig” for long-term time-lapse imaging of live adult zebrafish (**Fig 1**). The required equipment is compact and can easily be incorporated into any microscopy workspace with minimal disruption (**Fig 1A**). Intubation provides a continuous flow of oxygenated water through the gills of the fish (**Fig 1B-D**). The apparatus includes a custom-designed, 3-D printed plastic chamber (**Fig 1E,F**, **Supp. Files 1,2**) that incorporates a variety of useful features, including a digital thermometer monitoring water temperature and a single four-line peristaltic pump providing the necessary force for water flow through the single inflow and dual outflow tubes (**Fig 1B**), ensuring that inflow cannot exceed outflow (previous methods relied on use of multiple pumps, that could have differing flow rates or fail separately). The design also incorporates multiple additional safety features to protect expensive microscope systems including a water sensor (**Fig 1B,C**) connected to an emergency shut-off switch in case both of the redundant dual outflow tubes clog (**Fig 1B,C**), and an emergency overflow tube (**Fig 1B**) to shuttle excess liquid away and prevent chamber overflow if all else fails.

The schematic diagram in **Fig. 2A** provides an overview of the components of the intubation rig (**Supp. Table 1**) and how they are all assembled, while **Figs. 2B,C** show details of the specific arrangement of components in and connected to the imaging chamber. The commercially available chambered coverglass (Lab Tek II Imaging Dish, **Supp. Table 1**) is secured to the printed intubation chamber using silicone grease as shown in **Fig. 2D**. As shown in these schematics, the various parts are easily assembled to form the working unit without a great deal of technical knowledge. We have provided two different versions of the chamber design, one designed to fit on a regular Tokei Hit heated stage (**Fig 1E**), and one with four supports extending out from the base of the chamber body designed to fit in a 96 well plate holder on a stage designed to hold 96-well plates (**Fig 1F**). We designed the chamber to fit on the heated stages we had on our microscopes, but because we use a heat block to warm the water before it enters the chamber a heated stage is not required. Our design can be modified to fit other imaging systems using the computer aided design (CAD) software files that we provide (**intubation_chamber**.**stl, 96W_Intubation_chamber**.**stl**). The length of the four stage supports can be quickly modified using Google Sketchup or other widely available CAD software to fit on many different microscope stages from a variety microscope manufacturers. The designs can be 3-D printed using commercially available printing services (e.g., Xometry; **Supp. Table 1**). The rest of the intubation rig components including the tubing, water sensor, pumps, water bath, and thermometer are also commercially available (**Supp. Table 1**). This intubation rig is simple to replicate and cost effective.

To demonstrate the utility of this intubation set-up for high resolution long-term confocal imaging of living adult zebrafish, we performed time-lapse imaging of neutrophil recruitment to a wound site in an adult zebrafish with transgenically labeled neutrophils and lymphatic vessels. We made a small wound on the side of an adult *Tg(lyz:DsRed2)NZ50;Tg(mrc1a:eGFP)y251* double transgenic, *casper* mutant zebrafish. This line has red fluorescent neutrophils and green fluorescent lymphatic vessels (**Fig 3A**), and the use of the pigment-free *casper* (*roy orbison* and *nacre* double mutant) background provides improved tissue clarity for adult imaging (White et al., 2008). The endogenous blue autofluorescence of the scales allowed us to visualize where they were removed by wounding (**Fig 3B**). Over the course of nearly day-long time-lapse imaging of living adult zebrafish, we were able to image neutrophils moving into and accumulating in the wound area (**Fig 3C-J, Supp. Movie 1**). Interestingly, significantly increased neutrophil recruitment to the wound area doesn’t begin until two to three hours post injury (**Fig 3D,H, Supp. Movie 1**), which is much longer than an anesthetized zebrafish without intubation can survive, making intubation essential for studying this interesting process. We carried out three separate replicate overnight time-lapses (20-24 hours each), in each case observing robust neutrophil recruitment to the wound site. In some but not all cases we were able to fully revive adult fish after intubation (Supp Movie 2). Revival post intubation likely depends on the hardiness of the fish being intubated, and the “tightness” of the restraint being used. By using a 20X long-working distance objective, we were able to acquire high-magnification, high resolution images of highly active migratory neutrophils moving around and through lymphatic vessels close to the wound site (**Fig 3K-N, Supp Movie 1**). The extraordinary steadiness and reproducible positioning of the microscope field captured in these high-magnification images (**Supp Movie 1**) shows that animals intubated and mounted for imaging using this method are extremely stable and that long-term continuous imaging can be performed even at very high optical resolution. Together, our time-lapse images show that this intubation rig is useful both for large field of view imaging (**Fig 3C-J, Supp. Movie 1**) such as visualizing neutrophil trafficking and migration at large scale (**Supp Movie 1**) as well as for much higher magnification imaging for the visualization and tracking of the precise movements and morphologies of individual cells (**Fig 3K-N, Supp Movie 1**).

**Figure 3.**
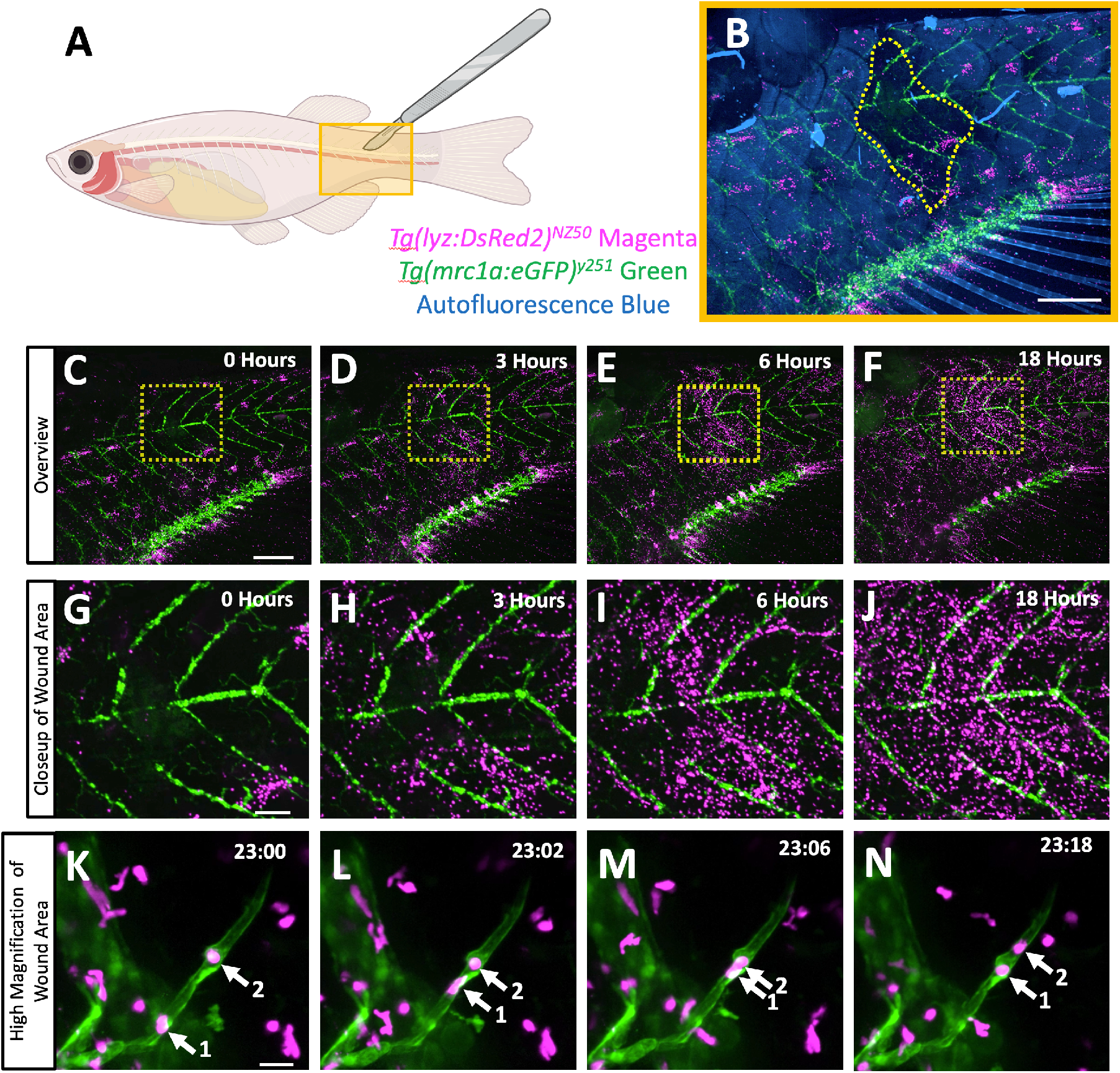
Long-term time-lapse imaging of neutrophil recruitment to a scale removal wound in an intubated adult zebrafish. **A**. Schematic diagram of an adult casper *Tg(lyz:DsRed2)*^*NZ50*^, *Tg(mrc1a:eGFP)*^*y251*^ double transgenic zebrafish with fluorescent neutrophils (magenta) and lymphatic vessels (green), as well as autofluorescent scales (blue). The approximate site of scale removal by abrasion with a scalpel is noted with a yellow box. **B**. An overview image of the wound area (yellow box in A) at the start of time-lapse imaging. The yellow dashed line notes the boundary of the site where autofluorescent blue scales were removed. **C-N**. Maximum intensity projection still images from long-term time-lapse confocal imaging of the adult fish in (B). **C-F**, Overview confocal images of the trunk at 0 (C), 3 (D), 6 (E), and 18 (F) hours. **G-J**. Close-up images of the boxed regions in panels C-F. **K-L**. High-magnification images of neutrophils (magenta) actively migrating in and around lymphatic vessels (green) in the recovering wound site of a live adult zebrafish after 23:00 (K), 23:02 (L), 23:06 (M), and 23:18 (N) of time-lapse imaging (hours:minutes). See **Supp. Movie 1** for the full time-lapse sequences including the images in panels C-N. Arrows note two neutrophils migrating inside a lymphatic vessel. Scale bars = 1 mm (B-F), 250 um (G-J), 25 um (K-N).

This study documents an easy to implement, easy to use, highly effective system for long-term time-lapse imaging of intubated adult zebrafish on an inverted microscope. This new system will make it possible to use high-resolution time-lapse optical imaging to study tissue regeneration in the fin, immune responses to injury, cancer metastasis, cellular trafficking through newly described intracranial lymphatic vessels, and other important processes taking place in adult zebrafish. In addition to being designed for widely used inverted microscope configurations, the design also incorporates many additional useful new attributes including precise temperature control, multiple water overflow safety prevention features, and an easily replicated and modified 3-D printed imaging chamber. The downloadable plans for this imaging chamber can easily be modified and adapted to create chambers to fit any commercial microscope stage. With ever-increasing use of the adult zebrafish as a model for studying disease, regeneration, and neurobiology, and other topics, this imaging “rig” will become an increasingly vital tool for long-term imaging of key events in adult fish.

## Supporting information

Chamber CAD file (change .txt to .stl to open)

Chamber CAD file (change .txt to .stl to open)

## ACKNOWLEDGEMENTS

The authors would like to thank members of the Weinstein laboratory for their critical comments on this manuscript. The Authors would also like to thank the Research Animal Branch of the *Eunice Kennedy Shriver* National Institute of Child Health and Human Development as well as the Charles River Staff for excellent animal care and husbandry.

## AUTHOR CONTRIBUTIONS

DC, BS, MVG, AVG and BMW designed experiments and developed the research ideas. DC performed the experiments. DC, BS, and BMW wrote the paper.

**Supplemental Table 1.** Materials List.

| Item | Description | Supplier | Part Number |
| --- | --- | --- | --- |
| 3D printed Chamber | Fits on microscope stage, holds LabTek dish | Xometry / Xometry.com | 3D renderings in supplement, Nylon 12 (SLS) |
| Lab Tek II Imaging Dish | Single chamber glass bottomed imaging dish #1.5 coverglass | ThermoFisher / ThermoFisher.com | Lab-TekII #155360 |
| Peristaltic Pump | Pumps fish water through chamber | World Precision Instruments / wpiinc.com | World Precision Instruments Peri-Star™ Pro Cat# PERIPRO -4L |
| Silicone Grease | Seals Lab Tek dish into chamber | Grainger / grainger.com | SG-ONE™ 24708 |
| Water supply tubing (pump) | Supplies water to chamber through peristaltic pump | US Plastics / usplastic.com | Tygon 3350 3/32" ID X 7/32" OD |
| Small inflow tubing | Brings water from supply tubing into chamber | US Plastics / usplastic.com | Tygon 3350 1/32" ID X 3/32" OD |
| Intubation Tubing | Small tube that fits in fish's mouth | World Precision Instruments / wpiinc.com | Flexible PE Tubing, 0.6mm ID, 1.1mm OD # 504280 |
| Overflow tubing | Large diameter tubing for emergency overflow | US Plastics / usplastic.com | Tygon S3 E-3603 |
| Outflow Elbows | Elbow that changes the direction of the outflow tube inside the 3D printed chamber | US Plastics / usplastic.com | 1/8" to 5/32" Kartell Polypropylene Elbow Connector # 64256 |
| Overflow Fitting | Attaches overflow tubing to chamber | US Plastics / usplastic.com | 1/2" to 5/8" Kartell Polypropylene Straight Hose Barb Connector #64114<br>(one end removed) |
| Overflow gasket | Gasket that holds overflow fitting in place | Grainger / grainger.com | Part of O-Ring Kit ASTM D2000 |
| Tube fitting kit | Attaches tubes together | World Precision Instruments / wpiinc.com | Luer-to-Tubing Coupler Assortment Kit # 504954 |
| Modeling Clay | Holds intubation tube in place | Various | Dixon Modeling Clay DIX00740 |
| Emergency Shut Off Switch | Modified "WasherWatcher Laundry Tub Overflow Protector" | HydroCheck, STAK Enterprises Inc. www.hydrocheckproducts.com | WasherWatcher Laundry Tub Overflow Protector |
| Sponge | Holds fish in place | Cole Parmer / Coleparmer.com | Jaece Identi-Plugs L800-D |
| Heat Block | Warms fish water | Southwest Science / southwestscience.com | SH100 Mini Dry Bath Hot Block, SWMINI-5 block insert |
| Digital Thermometer | Dual probe thermometer for monitoring incoming water temperature and chamber temperature | Fisher Scientific / Fishersci.com | Fisher Scientific Dual Thermometer 15-077-26 |
| Air Pump with Air stone | Aerates fish water in water bath | Various | Any, for small home aquarium |
| Tokei Hit Heated stage A | Keeps microscope samples warm | Tokei Hit / Nikon Instruments | Model: INUB-TIZB |
| Tokei Hit Heated stage B | Keeps microscope samples warm (fits 96 well plate) | Tokei Hit / Nikon Instruments | Model: STZF-TIZWX-SET |

**Supplemental Movie 1**

Time-lapse imaging of neutrophil recruitment to a scale removal wound in an intubated adult zebrafish. **0-4’’** Schematic diagram an adult casper *Tg(lyz:DsRed2)NZ50;Tg(mrc1a:eGFP)y251* double transgenic zebrafish with fluorescent neutrophils (magenta) and lymphatic vessels (green). The approximate site of scale removal by abrasion with a scalpel is noted with a yellow box. **5-10”** Adult fish being imaged in the intubation chamber on an inverted confocal microscope, using blue, green and purple (near-UV) excitation light. **11-31**” Time-lapse imaging of the wound site of an adult fish collected with a 2X objective from 0 – 19 hours post wounding and intubation **31-38”** Close-up of the wound site collected with a 10X objective at 19-20 hours post wounding and intubation. **38-44”** High magnification closeup of neutrophils (magenta) actively migrating in and around lymphatic vessels (green) in the recovering wound site of a live adult zebrafish collected with 20X objective from 23 – 24 hours post wounding and intubation.

**Supplemental Movie 2**

Video of an adult zebrafish swimming before and after a 20 hour intubation.

## SUPPLEMENTAL FILES

Supp CAD Files: intubation_chamber.stl, 96W_Intubation_chamber.stl

Supp Movie 1. intubation_neutrophil_movie1.mp4

Supp Movie 2. pre_and_post_intubation_movie2.mp4

